# Mapping of the Fondaparinux Binding Site of JR-FL gp120 by High Resolution Hydroxyl Radical Protein Footprinting and Computational Docking

**DOI:** 10.1101/207910

**Authors:** Sandeep K. Misra, Amika Sood, Paulo A. Soares, Vitor H. Pomin, Robert J. Woods, Joshua S. Sharp

## Abstract

The adhesion of HIV gp120 antigen to human cells is modulated in part by interactions with heparan sulfate. The HXB2 strain of gp120 has been shown to interact with heparin primarily through the V3 loop, although other domains including the C-terminal domain were also implicated. However, the JR-FL strain (representative of CCR5-interacting strains that make up newest infections) was shown to have a drastically lowered affinity to heparin due to the loss of several basic residues in the V3 loop, and deletion of the V3 loop in JR-FL gp120 was shown to abrogate some, but not all, heparin binding. Here, we use high resolution hydroxyl radical protein footprinting to measure the changes in protein surface oxidation levels that result from the binding of a model heparin fragment (fondaparinux). Protection in both the V3 loop and the N-terminus of JR-FP gp120 is observed. The well-defined composition of fondaparinux allowed us to perform docking simulations, which showed two clusters of fondaparinux binding: the V3 loop, and a domain consisting of the N- and C-termini. Together, the experimental and theoretical results indicate the heparin/heparan sulfate binding sites on JR-FL gp120 and the efficient interaction of fondaparinux, a widely exploited therapeutic carbohydrate, on gp120.

## Introduction

Human immunodeficiency virus (HIV) enters the CD4^+^ cells by interacting first with the primary receptor CD4 and then with a co-receptor, usually CXCR4 or CCR5 [1]. A glycoprotein, gp120, on the surface of HIV virus plays an important role in this process. Although gp120 consists of a polypeptide core of about 60 kDa, extensive *N*-linked glycosylation increases the molecular weight to approximately 120 kDa [2]. gp120 consists of five relatively conserved regions, C1 to C5, that fold into a core comprising two distinct domains termed “inner” and “outer”, and five variable regions, V1 to V5 [3]. A number of studies [4, 5–7] have shown that when gp120 binds to CD4, the gp120 core structure undergoes structural rearrangements, and this is more prominent in the inner domain. Upon binding of gp120 to CD4, the base of the V1/V2 region of the inner domain consisting of β2 and β3 strands is brought closer to a hairpin turn of the outer domain consisting of β20 and β21 strands and forms a four-stranded β-sheet located within the bridging sheet that connects the inner and the outer domain. This highly conserved structure forms the binding site for CCR5 or CXCR4 in conjunction with the V3 loop, [8, 9–11].

HIV virus can also infect CD4-cells by the interaction of gp120 with cell-surface associated heparan sulfate (HS) [12]. HS and the closely related heparin polysaccharide comprise a diverse family of heavily sulfated linear glycosaminoglycans (GAGs) that are covalently bound to proteins expressed on eukaryotic cell surfaces [13]. These GAGs have a variety of physiological and pathological functions, centered primarily on regulation of functional proteins. Signaling interactions occur between the cell surface GAGs and other cells or components of the extracellular matrix [14]. GAGs have been shown to play a variety of roles in HIV infection [4]. The binding of heparin/HS polysaccharides to gp120 has been attributed mainly to interactions with the V3 loop of gp120 in the HXB2 strain (which uses the CXCR4 co-receptor), although other regions of the protein have also been implicated. Four heparin-binding domains (HBDs) in the HXB2 strain have been identified, located in the V2 and V3 loops, in the C-terminal domain, and within the CD-4 induced bridging sheet [5]. While protein-GAG interactions are commonly mediated by basic amino acids, the exact identity of amino acids of gp120 involved in this interaction is not known, and a complex mixture of heparin structures was used, making the detailed structure-function analysis not very accurate [5]. Additionally, the identified HBDs contain several basic residues that are missing in the commonly studied JR-FL strain, which uses the CCR5 co-receptor and is more indicative of viruses most commonly found in early and chronic stages of HIV infection [15]. Previous work indicated that heparin bound poorly to JR-FL compared to HXB2, indicating that the mutation of these residues probably plays a strong role in modulating the interaction of gp120 with heparin/HS. Interestingly, the binding that was detected for JR-FL appeared to be partially mediated by the V3 loop, but deletion of the V3 loop did not completely reduce heparin binding on gp120 to background levels [16]. This suggests that JR-FL gp120 interacts with heparin through multiple domains, complicating any analysis of these interactions. A structural approach examining the interaction between a defined heparin oligosaccharide ligand and gp120 from the JR-FL strain would give more accurate information on the resultant complex, especially if investigated by both experimental and computational modeling techniques. To develop an understanding of the structural basis for these interactions, we employed a synthetic heparin oligosaccharide (fondaparinux) ligand and gp120 from the JR-FL strain in experimental hydroxyl radical protein footprinting (HRPF) analyses and computational docking studies that enabled us to fully define the binding modes.

Fondaparinux (Arixtra^®^) is a synthetic heparin pentasaccharide that has been previously used as a model heparin/HS oligosaccharide with defined structure (**Figure 1**), whose anticoagulant therapeutic activity derives from its ability to bind to antithrombin. Fondaparinux is a common oligosaccharide used in biophysical studies of GAG interactions with proteins, in which a readily available GAG oligosaccharide of the defined structure is required for the study [17, 18–20]. The synthetic pentasaccharide includes a 3-O-sulfated glucosamine, which has been implicated previously in adhesion of some viruses to cell surfaces [21, 22]. The experimental characterization and computational modeling of the contact sites between gp120 and fondaparinux will provide a better understanding of how gp120 interacts with fondaparinux and suggest possible methods for blocking the entry of the HIV virus into the host cells.

**Figure 1.**
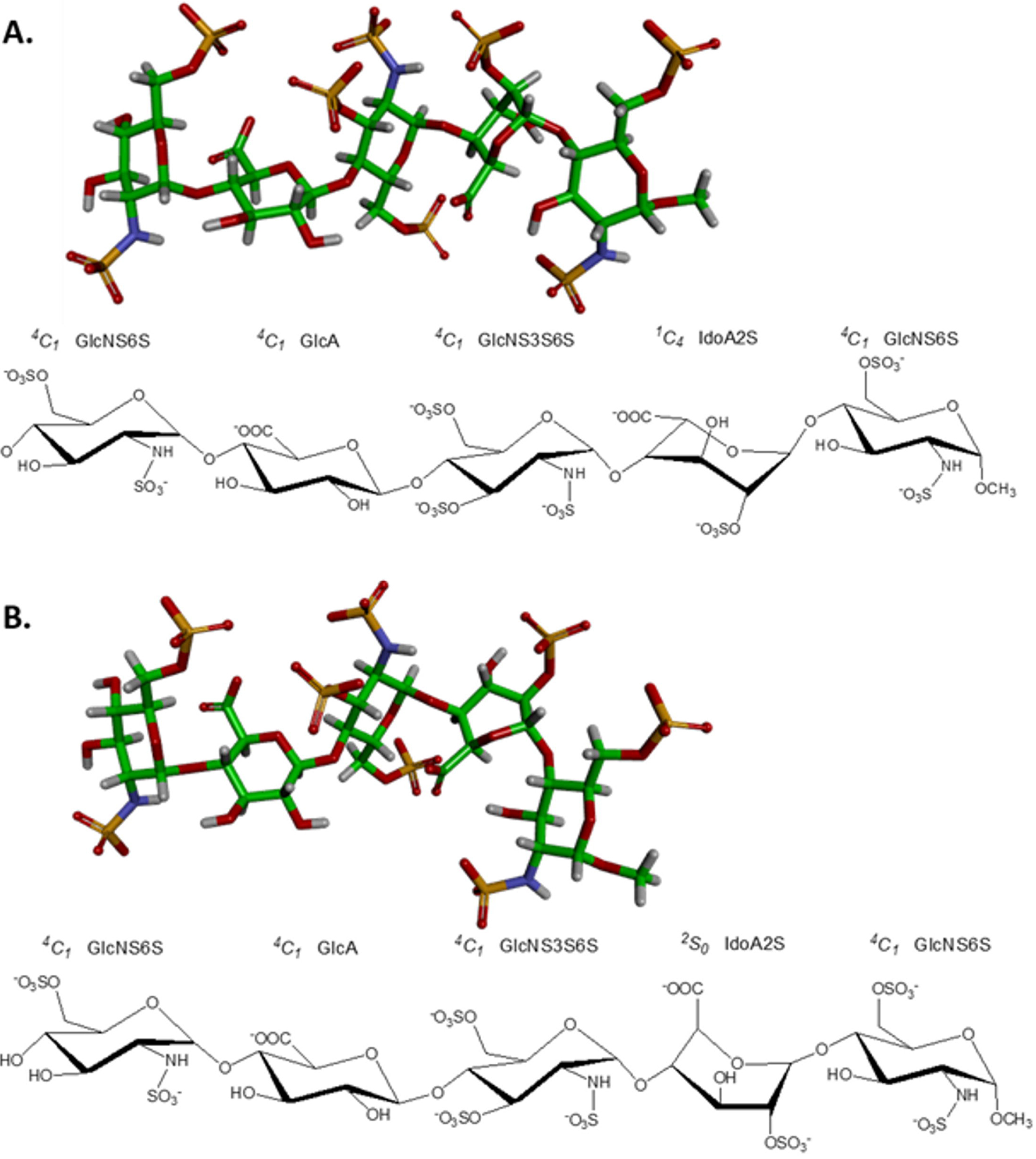
Structure of fondaparinux with the iduronic acid in the ^1^C_4_ (A) or ^2^S_0_ (B) configurations. GlcNS6S, GlcA, GlcNS3S6S and IdoA2S stand respectively for 2N-, 6O-disulfated glucosamine, glucuronic acid, 2N-, 3,6O-trisulfated glucosamine and 2O-sulfated iduronic acid.

In order to map the fondaparinux binding region of gp120, we performed high-resolution hydroxyl radical protein footprinting (HR-HRPF) by fast photochemical oxidation of proteins (FPOP). HR-HRPF is a relatively recently developed method designed to probe changes in the topography of a protein [23, 24–33]. HR-HRPF by FPOP stably modifies the solvent accessible amino acid side chains without unfolding or deforming the protein conformation during the time scale of modification, allowing heavy surface labeling of the native structure [34]. To measure changes in HR-HRPF reactivity, the stable modifications to the protein side chains are analyzed by liquid chromatography (LC) and mass spectrometry (MS) techniques, and the levels of reactivity are relatively quantified by comparison of the modified and unmodified peptides. A number of studies have employed MS-based HRPF for investigating protein conformational changes as well as protein-protein interactions, protein folding and protein-ligand interactions [35, 37–43].

Some recent works from our group have established that using electron transfer dissociation (ETD)-based methods for quantifying multiple adjacent sites of isomeric oxidation products allows accurate quantification of changes in the HRPF down to single-amino acid spatial resolution providing structural information with higher spatial resolution and more accuracy [43, 44–45]. Specific poses of the molecular interactions between a defined heparin oligosaccharide and gp120 can be developed by examining changes in the HR-HRPF of gp120 in the presence and in the absence of fondaparinux and comparing the generated results with computational docking simulations using a recently released model of the fully glycosylated JR-FL gp120 [43].

## Materials and Methods

### Materials and Reagents

Purified JR-FL gp120 was purchased from Immune Technology Corp (New York, USA). Fondaparinux, purchased from GlaxoSmithKline (Brentford, UK), was injected onto a Bio-Gel P-2 (Bio-Rad, USA) size exclusion column and eluted with milli-Q water at a flow rate of 1 mL/7.5 min for desalting and further purification. Fractions of 20 mL were collected and assayed by metachromasia using 1 mL of 1,9-dimethylmethylene blue (DMB). Fractions of 3 μL were collected and tested for the presence of NaCl using 200 μL of AgNO3. All fractions that were positive for DMB and negative for AgNO3 were pooled and lyophilized. Catalase and dithiothreitol (DTT) were obtained from Sigma-Aldrich (St. Louis, MO, USA). Sodium phosphate, LC-MS grade formic acid and hydrogen peroxide were purchased from Fisher Chemicals (Fair Lawn, NJ, USA). Adenine and L-glutamine were purchased from Acros Biologicals. Methionine amide was obtained from Bachem (Torrance, CA, USA). Sequencing-grade modified trypsin was purchased from Promega Corp. (Madison, WI). PNGase F (500000 units/mL) was purchased from New England Biolabs (Ipswich, MA). All reagents were used without further purification. Purified water (18.2 MΩ) was obtained from a Synergy UV system (Millipore, Billerica, MA).

### FPOP of gp120-Fondaparinux

FPOP was performed as previously described [33]. Briefly, a final concentration of 2 μM gp120 JR-FL protein was incubated with 20 mM of phosphate buffer, pH 7.4 either with or without 2 μM fondaparinux at room temperature for 1 h containing 17 mM glutamine. This mixture contained 1 mM adenine as a radical dosimeter to monitor the available radical dose in each sample [45]. 2 μL of freshly prepared 1M hydrogen peroxide was added to the sample mixture immediately prior to laser irradiation. The total 20 μl of this sample mixture irradiated by flow through the path of the pulsed ultraviolet laser beam, with each volume illuminated by no more than a single pulse of the laser. Briefly, a Compex Pro 102 KrF excimer laser (Coherent, Germany) was run at ~9.0 mJ/mm^2^/pulse, with a laser repetition rate of 15 Hz. The flow rate was adjusted to 13.2 μL/min to ensure a 20% exclusion volume between irradiated segments to help to accountfor sample diffusion and laminar flow [46]. After illumination, each replicate was collected in a microcentrifuge tube containing 25 μL of quench mixture that contained 0.5 μg/μl H-Met-NH2 and 0.5 μg/μl catalase to eliminate secondary oxidants such as remaining hydrogen peroxide, protein peroxides, superoxide, etc. The samples were incubated in the quench solution for 30 min at room temperature with pipet mixing. Control samples were handled in the same manner as samples with FPOP, except they were not laser irradiated. All experiments were performed in triplicate for statistical analysis.

Following laser irradiation and quenching, 2 μL of the sample mixture was analyzed at 260 nm on Nano-Drop 2000c UV-vis spectrophotometer (Thermo Scientific). This step was performed to ensure that each sample received a comparable effective dose of hydroxyl radicals. The amounts of adenine oxidation were similar after HR-HRPF and measured by UV among all tested samples with an average of around 30% oxidized adenine indicating that all samples received the same level of hydroxyl radicals.

### Sample Preparation

50 mM Tris, pH 8.0 and 1 mM CaCl_2_ was added to the protein samples after FPOP. 5 mM of dithiothreitol was added and incubated the mixture at 80°C for 20 minutes to denature and reduce the protein. After the mixture had been cooled to room temperature, a 1:20 trypsin weight ratio was added to the samples for overnight digestion at 37 °C with sample rotation. Digestion was terminated by adding 10 mM DTT and heating the samples to 95 °C for 10 min. Finally, when the samples cooled to room temperature, 150 units of PNGase F were added to the digested samples and the samples were incubated at 37 °C for 16 h. The reaction was terminated by adding DTT to a final concentration of 20 mM and heating the sample to 95 °C for 10 min. Samples were stored at −20 °C until analysis was performed on nano-LC-MS/MS system.

### Mass Spectrometry

The protein samples were analyzed on an Orbitrap Fusion instrument (Thermo Fisher Scientific) controlled with Xcalibur version 2.0.7 (Thermo Fisher, San Jose, CA). Samples were loaded onto an Acclaim PepMap 100 C18 nanocolumn (0.75 mm × 150 mm, 2 μm, Thermo Fisher Scientific). Separation of peptides on the chromatographic system was performed using mobile phase A (0.1% formic acid in water) and mobile phase B (0.1% formic acid in acetonitrile) at a rate of 300 μL/min. The peptides were eluted with a gradient consisting of 2 to 35% solvent B over 25 min, ramped to 95 % B over 3 min, held for 3 min, and then returned to 2% B over 3 min and held for 8 min. Peptides were eluted directly into the nanospray source of an Orbitrap Fusion instrument using a conductive nanospray emitter obtained from Thermo Scientific. All data were acquired in positive ion mode. Both collision-induced dissociation (CID) and ETD were used to fragment peptides, with an isolation width of 3 *m/z* units. The spray voltage was set to 2300 volts, and the temperature of the heated capillary was set to 300 °C. In CID mode, full MS scans were acquired from m/z 200 to 2000 followed by eight subsequent MS2 scans on the top eight most abundant peptide ions. In ETD mode, the parent ions of all identified peptides were included in the parent mass list. ETD-based precursor activation was performed for 100 ms, including charge state-dependent with supplemental activation of 5V CID energy.

### HR-HRPF Data Analysis

Unoxidized gp120 controls, oxidized gp120, and oxidized complex of gp120-fondaparinux complex peptide sequences were initially identified using ByOnic version v2.10.5 (Protein Metrics) using gp120 JR-FL sequence database. The enzyme specificity was set to cleave the protein after lysine and arginine. Deamination on both asparagine and glutamine and all possible major oxidation modifications [47] were included as variable modifications for database searches. All tandem mass spectral assignments and sites of oxidation were verified manually. The peak intensities of the unoxidized peptides and their corresponding oxidation products observed in LC-MS were used to calculate the average oxidation events per peptide in the sample. Peptide level oxidation was calculated by adding the ion intensities of all the oxidized peptides multiplied by the number of oxidation events required for the mass shift (e.g., one event for +16, two events for +32) and then divided by the sum of the ion intensities of all unoxidized and oxidized peptide masses as represented by equation 1.

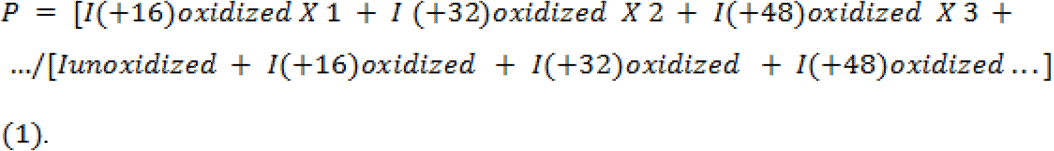

where *P* denotes the oxidation events at the peptide level and *I* values are the peak intensities of oxidized and unoxidized peptides. gp120 control samples that were not exposed to laser irradiation, but identical to the experimental samples otherwise, were analyzed to ensure that background oxidation would not be high enough to interfere with HR-HRPF data. These control samples showed <8% of the oxidation extent of the gp120 sample (data not shown). The interference of this small background is not significant enough to affect the comparison of the oxidation extent between gp120 and gp120-fondaparinux samples.

In cases in which oxidation at specific sites can be identified on the basis of either the mass differences between nonisomeric oxidation products or the presence of only a single oxidation site within a peptide as confirmed by CID and/or ETD, residue level quantitation is calculated from the LC-MS signal intensities of each peptide containing a specific oxidized amino acid (I_oxidized_), relative to the total of all intensities associated with that peptide sequence (I_oxidized_ +I_unoxidized_) according to equation2:

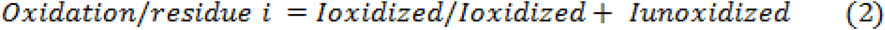

In cases where oxidation of a peptide results in a mixture of isomers with oxidation at different sites, residue level quantitation is calculated from the fragment ion intensities from ETD to determine the oxidation extent at a specific residue site based on our previous studies [33, 43]. Briefly, an oxidized peptide with multiple sites of oxidation can generate both oxidized and unoxidized sequence ions in its tandem mass spectrum. The fractional oxidation of a given sequence ion is defined as the ratio between the oxidized sequence ion intensity and the sum of the intensity of the corresponding oxidized and unoxidized sequence ion as represented in equation 3:

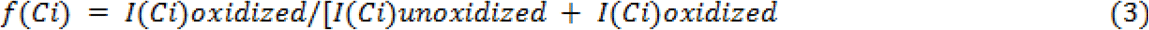

Here, *f*(*Ci*) denotes the fractional oxidization of sequence ion *I* (e.g., oxidized C3 ions generated by ETD) and *I*(*Ci*) denotes the intensity of sequence ion *i*, whether it is the oxidized or unoxidized form. The absolute level of oxidation for a given amino acid residue *i* is based on both the average oxidation event of the peptide and the fractional oxidation of the corresponding sequence ions, as shown in equation 4:

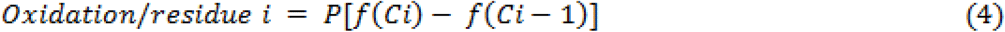

where *P* is the average oxidation events per peptide as derived from eq 1 and the term in brackets is the fractional oxidation difference of two adjacent sequence ions *i* and *i* − 1. The multiplication of the average number of oxidations per peptide by the fraction of that oxidation that occurs on a given amino acid residue yields the average oxidation events per residue. The protection by fondaparinux binding is defined as the ratio of the difference in oxidation extent between the gp120 sample alone and the gp120-fondaparinux binding sample to the oxidation extent of the gp120 sample alone. This is shown in eq 5:

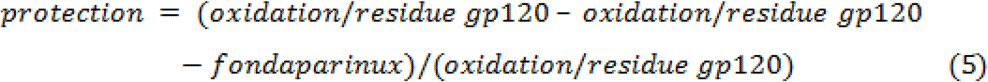

### gp120-Fondaparinux Complex Docking

The glycosylated model of gp120 previously published [43] was modified by the addition of the terminal structure from a previously published crystal structure of JR-FL SOSIP.664 (PDB ID: 5FYK) [48]. The termini were minimized *in vacuo* by steepest descent minimization for 5000 steps followed by 20000 steps of conjugate gradient minimization with the sander.MPI version of AMBER14 [49] using the Amber ff14SB force field. The resulting model was used for docking. The three-dimensional structures of fondaparinux, one with IdoA in ^1^C_4_ chair configuration and the other with IdoA in ^2^S_0_ skewed-boat configuration were generated using GLYCAM-Web server (http://www.glycam.org).

Docking was performed using Autodock 4.2.6 (AD4.2.6) [50] and Autodock Vina 1.1.2 (ADV) [51]. All input files were prepared for docking using AutoDock Tools 1.5.6 (ADT) [50]. For blind docking (entire protein surface used as the search space) with AD4.2.6, Gasteiger charges [52] were assigned to prepare protein and ligands and a grid box with 1Å spacing was employed for all runs, centered at the gp120 protein center. 100 runs of the Lamarckian Genetic Algorithm were performed, with 2,500,000 energy evaluations per run, and a population size of 200. For targeted docking with ADV, the exhaustiveness value was set to 80, while all the other parameters were set to their default values. Both the protein and ligand remain rigid for docking in all cases. VMD [53] was used to calculate the average structure, the center of the structure and for image rendering.

Solvent accessibility surface area (SASA) was calculated using NACCESS [54] program. Per-residue relative SASA was used for the analysis. Docking poses representing the lowest energy configurations of the iduronic acid in either the ^1^C_4_ or ^2^S_0_ configurations at the two docking cluster sites were shown using PyMol.

## Results

### HR-HRPF of JR-FL gp120 and the JR-FL gp120-Fondaparinux Complex

HR-HRPF topographical analysis of gp120 was performed both in the absence and presence of equimolar concentrations of fondaparinux. After FPOP experiments, we obtained 29 peptides identified by trypsin digestion of the gp120 covering 83.3% of the protein sequence as presented in **Figure 2**. Of these, 21 peptides were observed oxidized by the FPOP oxidative labeling with either +16, +32 or +48 Da mass shift. All the peptide identifications and the site(s) of oxidation were manually validated by CID and/or ETD MS/MS. A histogram showing the amount of oxidation of each peptide in the presence and absence of fondaparinux is shown in **Figure 3**.

**Figure 2.**
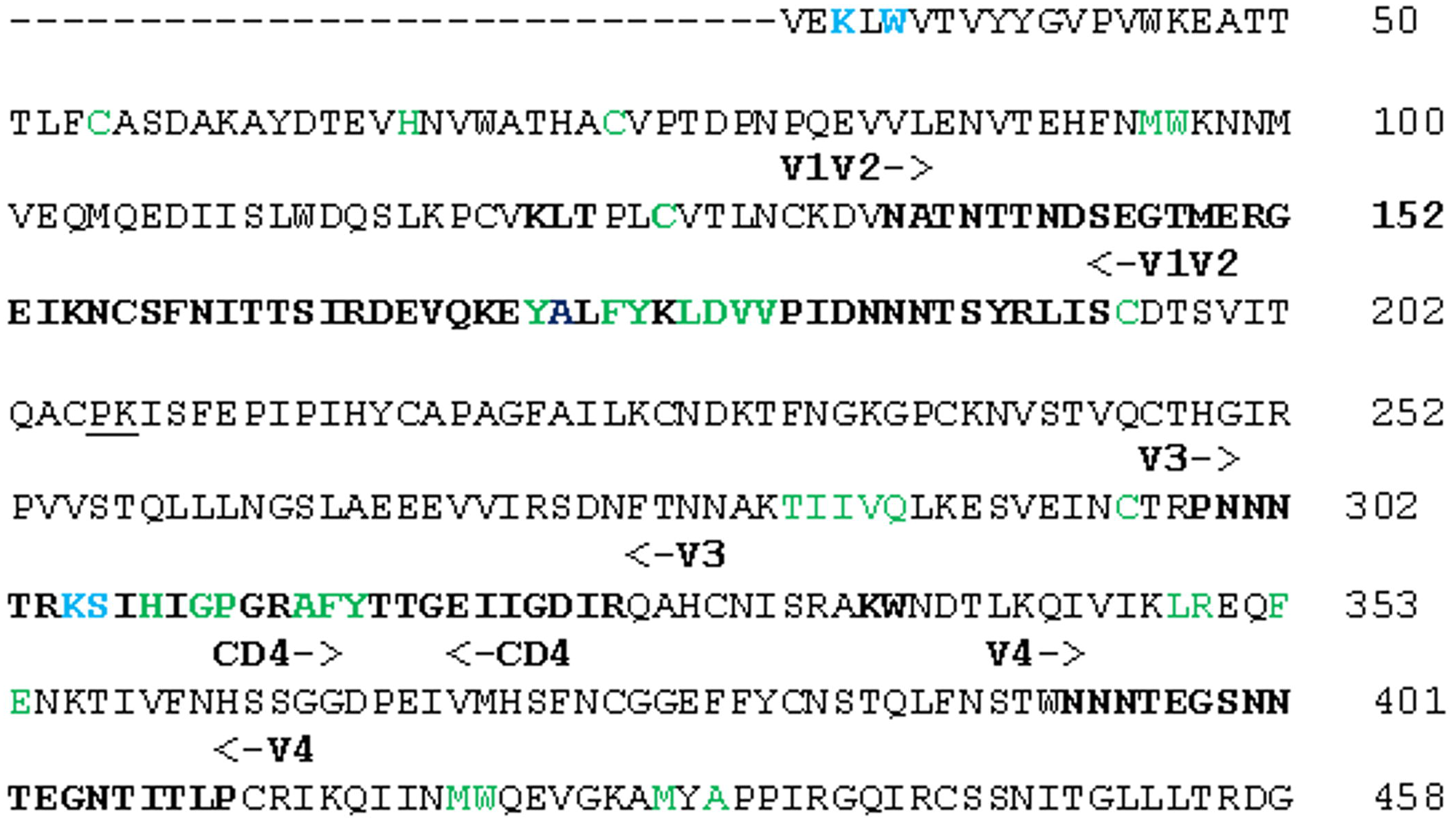
The sequence of HIV-1 gp120 (JR-FL) numbered using the HXB2 convention. Sites of oxidation that show statistically significant change in topography are shown in green; sites of oxidation that show a statistically significant decrease in solvent accessibility upon fondaparinux binding (*p* < 0.05) are shown in blue. The variable domains are labeled above the sequence.

**Figure 3.**
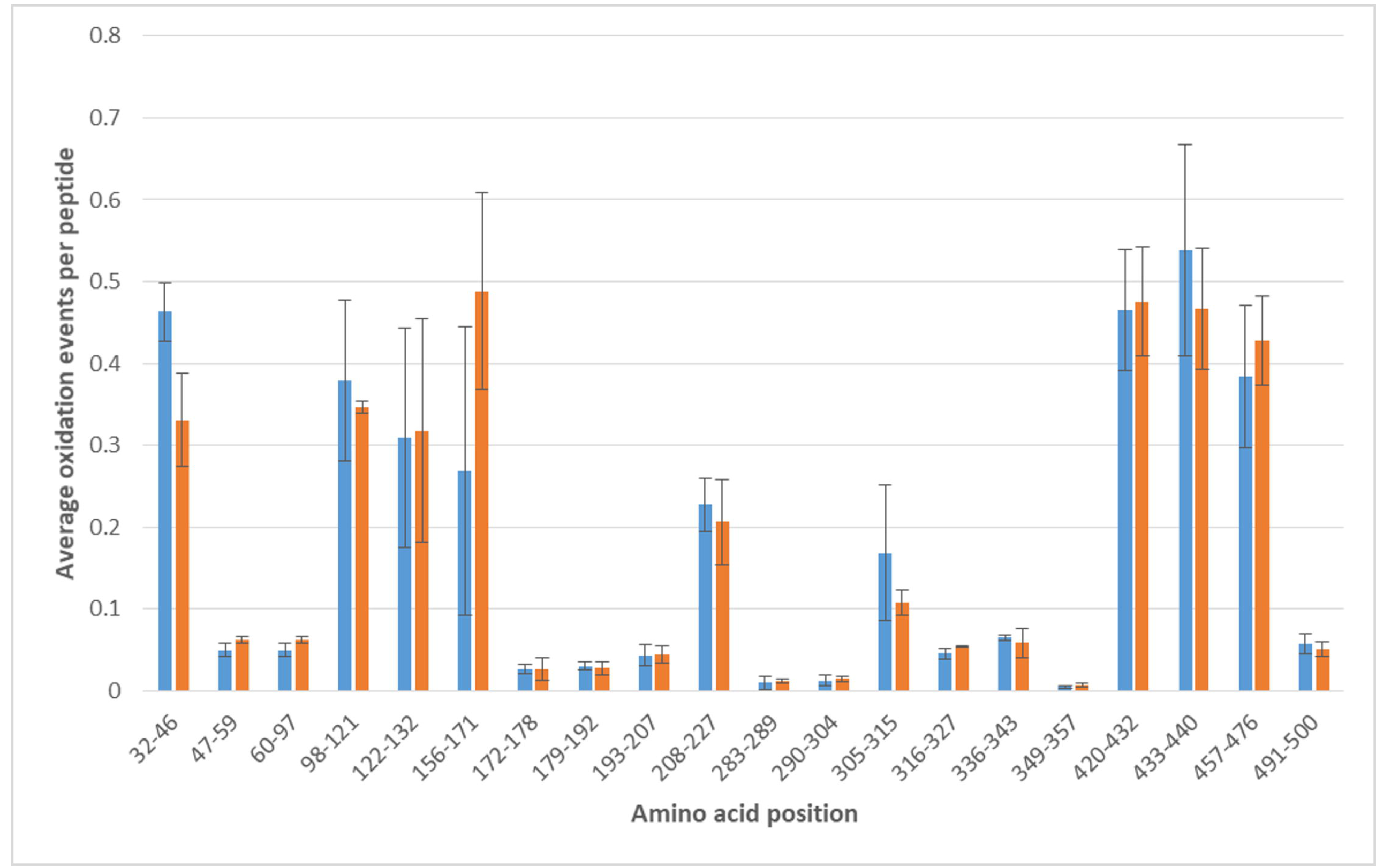
Peptide level HRPF of JR-FL gp120 in the absence (blue) and presence (orange) of equimolar fondaparinux. Error bars represent one standard deviation from a triplicate measurement.

Using ETD-based MS/MS data, high spatial resolution HR-HRPF measurements of the level of oxidation of each amino acid or small peptide fragment were made in the presence and absence of equimolar fondaparinux as described in Materials and Methods. Changes in the amount of oxidation of each measured amino acid due to fondaparinux binding are shown in a volcano plot in **Figure 4**. The amino acids K32, W35, and the K305-S306 dipeptide fragment showed protection upon fondaparinux binding that was both statistically significant (*p* < 0.05) and of substantial magnitude (> a 30% decrease in oxidation). No other amino acids showed changes in the HR-HRPF footprint that even approached statistical significance.

**Figure 4.**
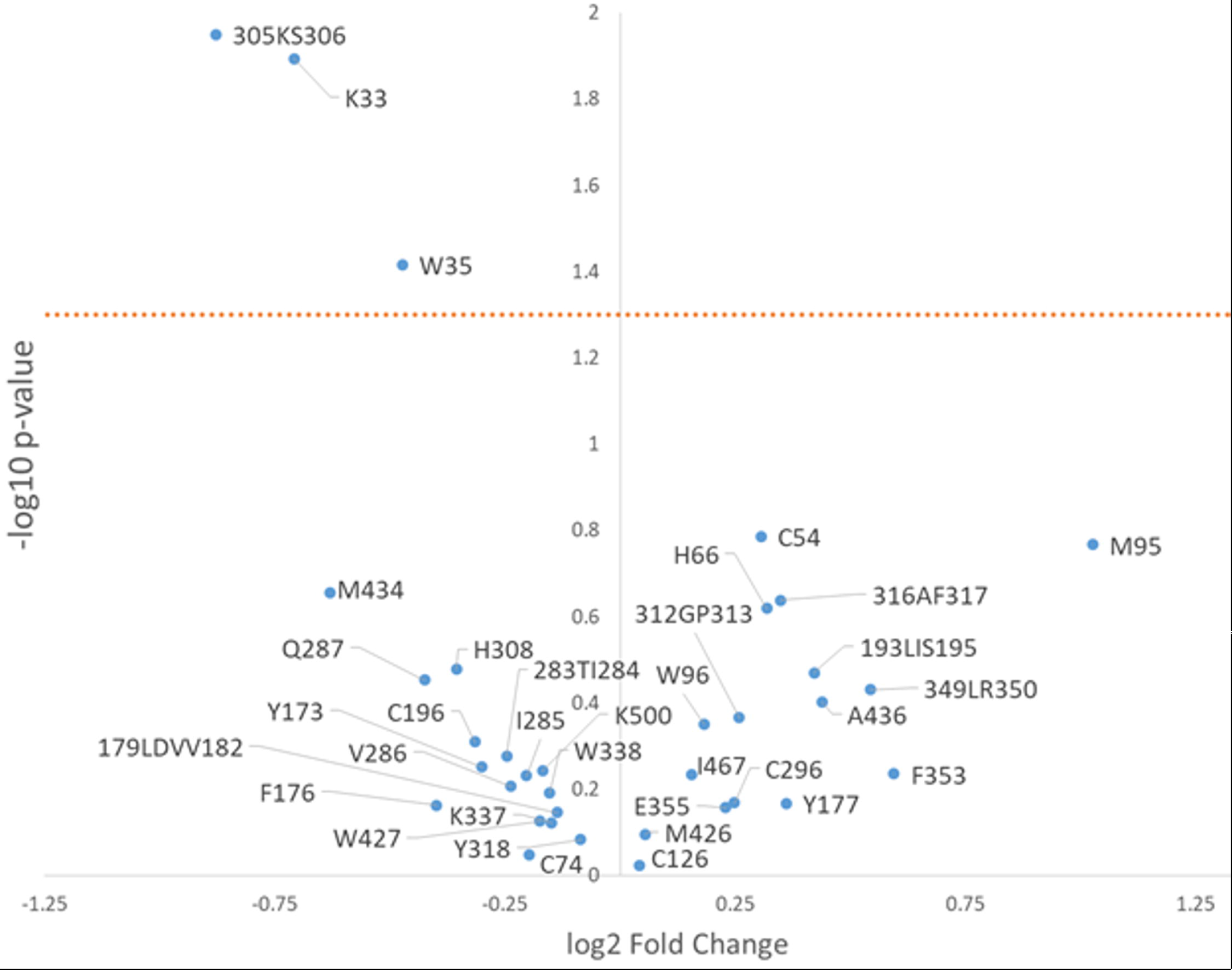
Volcano plot of the HR-HRPF data from JR-FL gp120 in the presence and absence of fondaparinux. A fractional fold change (i.e. a negative log2 fold change) represents a decrease in oxidation (and therefore solvent accessible surface area) upon binding to fondaparinux. The *p*-value was determined by a two-tailed Student’s t-test from triplicate samples. The orange dotted line represents *p* = 0.05.

### Docking of fondaparinux with JR-FL gp120

Previous work from our group generated a glycosylated model of JR-FL gp120 [43]. However, this model lacked the N- and C-termini of the JR-FL structure, due to their absence in the template structure. Given the protection of amino acids at the extreme N-terminus of JR-FL gp120 upon fondaparinux binding, the N- and C-termini modeled from a recent X-ray crystal structure of JR-FL SOSIP.664 (PDB ID: 5FYK) [48] were added to our previously published model and minimized. Fondaparinux was docked to this model in two separate blind docking runs using AD4.2.6: one with the iduronic acid in the ^1^C_4_ chair conformation and one with it in the ^2^S_0_ skew-boat form. In both the cases, the docked structures clustered on either the electropositive patch close to the V3 region of gp120 (site 1) or at the patch formed by both the N- and C-terminal region (site 2) (**Figure 5**). Targeted docking was then performed using ADV, by placing a grid box of size 30 × 30 × 30Å at the center of the cluster from the previous step (**Figure 6**).

**Figure 5.**
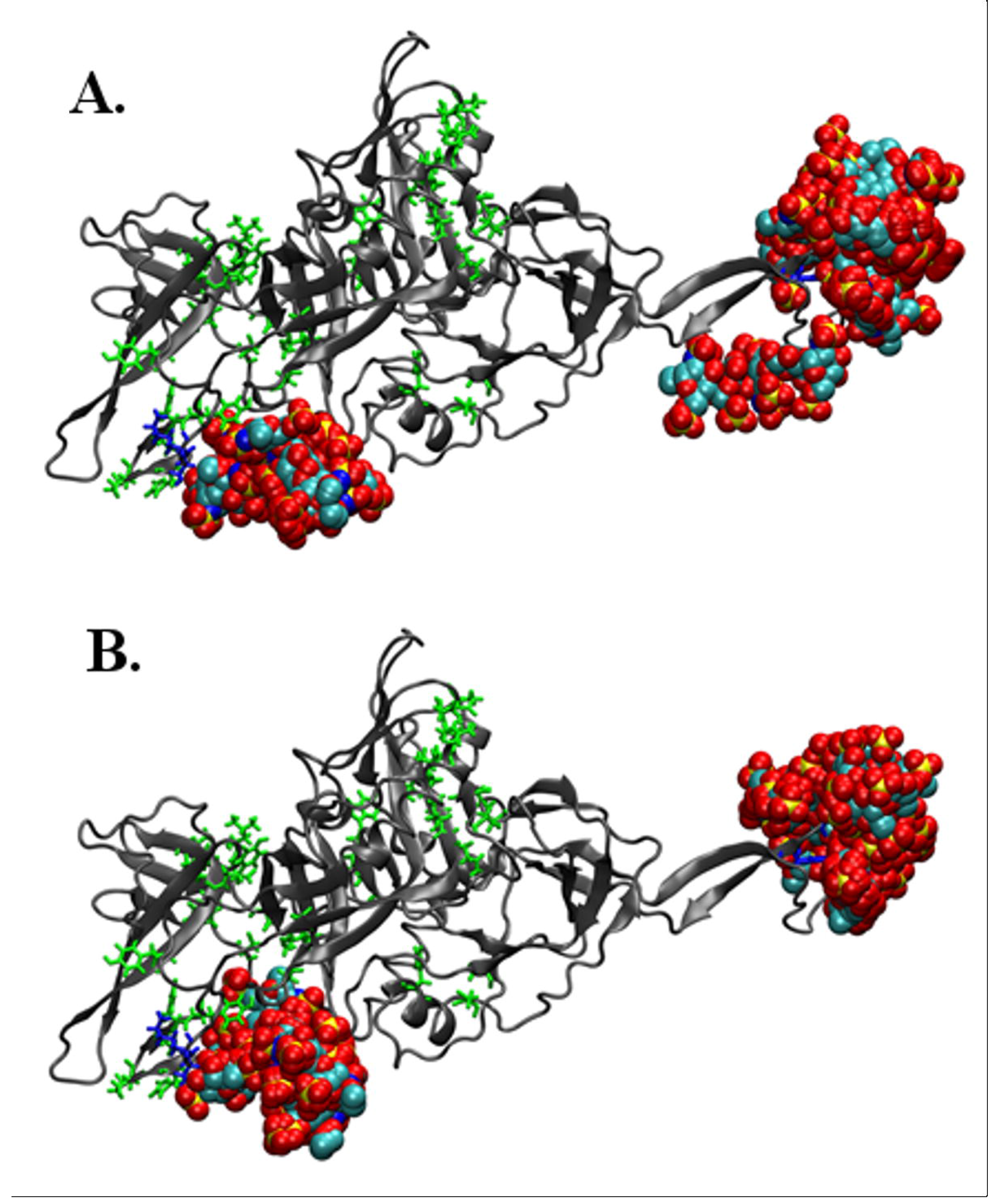
Blind docking of fondaparinux with glycosylated JR-FL gp120 using Autodock4.2.6. Two clusters were formed, one near the V3 loop, and the other near the N- and C-termini. Two ring configurations for the iduronic acid of fondaparinux were used: the ^1^C_4_ chair conformation (**A**) and the ^2^S_0_ skew-boat conformation (**B**). Oxidized amino acid side chains in HR-HRPF are shown as stick representations. Amino acids that showed statistically significant protection are colored blue, and amino acids that showed no significant protection are shown in green. While the N-linked glycans are present in the docking process, they have been removed from this representation for clarity.

**Figure 6.**
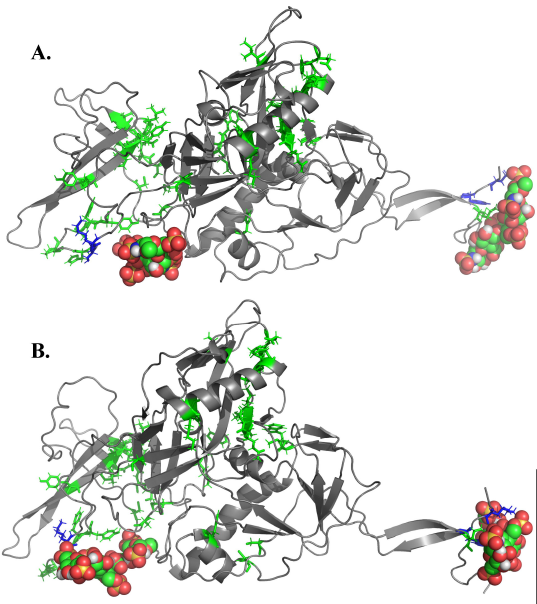
Computational docking of glycosylated JR-FL gp120 with fondaparinux with the iduronic acid ring conformation in either the ^1^C_4_ chair conformation (A.) or the ^2^S_0_ skew-boat conformation (B.) Fondaparinux is shown as ball models, and amino acids oxidized in HR-HRPF experiments are shown as stick models green: no protection upon fondaparinux binding; blue: significant protection upon fondaparinux binding. Blind computational docking using AD4.2.6 gave similarly plausible poses clustered in two regions: the V3 loop near the 305-KS-306 amino acids (lower left) and the N- and C-terminal region near K33 and W35 (far right). The top pose from targeted docking using ADV from each cluster are shown. While the N-linked glycans are present in the docking process, they have been removed from this representation for clarity.

Ligand binding shields protein residues from interacting with the solvent. Therefore, this shielding can be used to specify the protein region involved in docking. These data also reflect the experimental changes that are measured by HR-HRPF [55, 56]. Changes in solvent accessibility were estimated using the difference between the SASA values of unbound and docked protein structures (ΔSASA). Average ΔSASA was calculated using all the 20 docked poses in each case. Amino acids showing protection upon fondaparinux binding of at least 10 Å^2^ for each fondaparinux IdoA ring conformer and each binding site cluster are shown in **Table 1**.

## Discussion

The N-terminal amino acids within JR-FL gp120, K33 and W35, that exhibit protection upon fondaparinux binding (*p* < 0.05) suggest either a direct binding event of fondaparinux at the N-terminal domain or an allosteric conformational change upon binding at the V3 loop revealed by the protection of the 305-306 KS dipeptide. The lack of change in oxidation in the many probed sites within gp120 argues against a widespread allosteric change upon binding. The fact that blind docking simulations of fondaparinux (in either the ^1^C_4_ or ^2^S_0_ ring conformations of the iduronic acid) to glycosylated JR-FL gp120 cluster to the precise two regions identified by HR-HRPF suggest that these are direct binding events being probed.

While the docking results agree with the HR-HRPF data on the general location of binding, the agreement is not perfect with the observed HR-HRPF protection. At the V3 loop binding site, the 305-306 KS dipeptide exhibited low levels of protection for both the ^1^C_4_ and ^2^S_0_ iduronic acid ring conformations (~1.3 Å^2^ for K305 and ~7.2 Å^2^ for S306). Examination of the docked models at this site shows the rotation of the K305 side chain away from the site of binding to form an ionic interaction with E172 (**Figure 7**). This ionic interaction does not appear to be an artifact of the molecular dynamics simulation; it is also present in the recent X-ray crystal structure of the JR-FL SOSIP.664 Env trimer [48]. One of the limitations of the docking approach with the fully glycosylated protein of this size is the need to hold the protein static in order to keep docking times reasonable. K305 is present in a flexible region of the V3 loop. Based on the HR-HRPF data and the docking models, it is a reasonable hypothesis that this lysine swings out in the presence of fondaparinux to form a salt bridge with the 6-O-sulfate of the internal GlcNS3S6S and/or the N-sulfate of the GlcNS6S on the methylated reducing end of fondaparinux.

**Figure 7.**
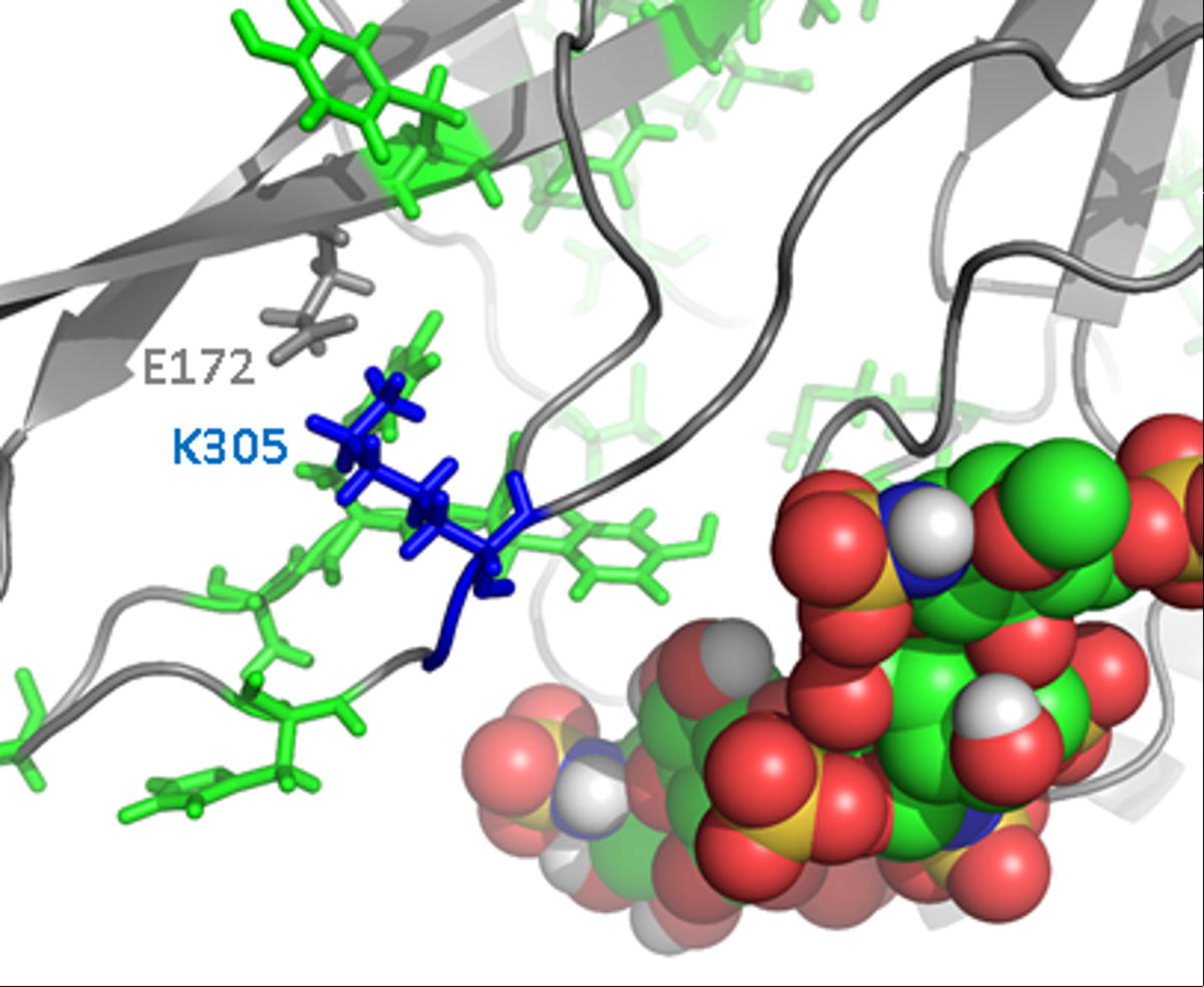
Zoomed in representation of K305 with fondaparinux (^1^C_4_) docked in the V3 loop site. Rotation of K305 away from E172 could put the amine in proximity to two anionic sulfates of fondaparinux (sulfur – orange; oxygen – red).

**Table 1.** **Amino acids protected ≥ 10Å^2^ by fondaparinux binding in each docking cluster**

| V3 loop Cluster |  |  |  | N-Terminus Cluster |  |  |  |
| --- | --- | --- | --- | --- | --- | --- | --- |
| <sup>1</sup>C<sub>4</sub> |  | <sup>2</sup>S<sub>0</sub> |  | <sup>1</sup>C<sub>4</sub> |  | <sup>2</sup>S<sub>0</sub> |  |
| Residue | $\Delta\text{SASA}$<br>($\text{\AA}^2$)<sup>†</sup> | Residue | $\Delta\text{SASA}$<br>($\text{\AA}^2$)<sup>†</sup> | Residue | $\Delta\text{SASA}$<br>($\text{\AA}^2$)<sup>†</sup> | Residue | $\Delta\text{SASA}$<br>($\text{\AA}^2$)<sup>†</sup> |
| T63 | 10 ± 14 | K121 | 22 ± 14 | V31 | 17 ± 21 | V31 | 11 ± 17 |
| K121 | 17 ± 15 | P124 | 24 ± 20 | E32 | 21 ± 8 | E32 | 13 ± 9 |
| P124 | 13 ± 15 | C205 | 11 ± 5 | **K33** | **31 ± 11\*** | **K33** | **27 ± 14\*** |
| P206 | 35 ± 21 | P206 | 39 ± 19 | **W35** | **15 ± 9\*** | **W35** | **21 ± 8\*** |
| T303 | 13 ± 20 | R304 | 18 ± 12 | V36 | 10 ± 21 | V36 | 15 ± 22 |
| R304 | 17 ± 12 | I307 | 31 ± 18 | **K500** | **21 ± 13\*** | P498 | 10 ± 15 |
| I307 | 19 ± 14 | **Y318** | **14 ± 12\*** | K502 | 19 ± 10 | **K500** | **15 ± 13\*** |
| P438 | 10 ± 7 | P437 | 12 ± 6 | R504 | 13 ± 14 | K502 | 13 ± 11 |
| R440 | 23 ± 13 | P438 | 11 ± 6 | V506 | 31 ± 16 | V506 | 34 ± 24 |
|  |  | R440 | 16 ± 9 |  |  |  |  |

<sup>†</sup>Average $\Delta\text{SASA}$ values from a cluster of twenty structures, ± one standard deviation \*HR-HRPF oxidized residues are colored: green – no protection upon fondaparinux binding; blue – protection upon fondaparinux binding.

As mentioned by Lortat-Jacob and co-workers, JR-FL gp120 is missing several basic residues in the V3 loop that are thought to drive binding of heparin to this domain in CXCR4-binding strains, and as a result, has a much lower binding affinity that is only partially dictated by the V3 loop [16]. We hypothesize that it is the reduction of affinity in this high-affinity binding site that causes a segregation of fondaparinux to both the V3 loop and the terminal heparin binding domain in JR-FL, as the sequence is almost fully conserved between HXB2 and JR-FL strains in the terminal regions involved, the only exception being T31 in HXB2 replaced by V31 in JR-FL. This terminal binding domain is probably inclusive of the C-terminal HBD described for HXB2 gp120, as the N-terminus detected by HR-HRPF here as a binding site for fondaparinux forms a two-strand anti-parallel beta-sheet with the C-terminus, holding the residues identified as protected in the N-terminus here by HR-HRPF in direct proximity to the C-terminal HBD3 identified by Lortat-Jacob and coworkers [5]. We further hypothesize that this terminal domain is the major contributor to heparin binding above background in JR-FL with the V3 loop deleted [16].

It should be noted that previous work on the interaction of full-length heparin with Robo1 found two distal sites of heparin binding, one at an internal flexible region and one at the N-terminus. Mutation of either of the two binding sites decreased the affinity of Robo1 to heparin by a similar amount, illustrating the role of avidity in the binding of these long polysaccharides to multiple binding sites of a single protein [33]. We hypothesize that these two distal binding sites may play a similar role in gp120, where tethering of the gp120 to a heparin/HS polysaccharide at one of the two gp120 heparin binding sites increases the binding at the second binding site, and engagement of both binding sites increases the functional affinity of gp120 on the cell surface polysaccharide. HR-HRPF of JR-FL gp120 complexed with full-length heparin may be informative; however, computational modeling of such a system becomes less useful due to the heterogeneity and polydispersity of heparin, and the size of the gp120-heparin complexes.

Our data, supported by blind docking, indicate that fondaparinux in an equimolar ratio can bind to JR-FL gp120 at low micromolar concentrations in sufficient amounts to detect using HR-HRPF measurements of changes in the protein topography. It remains to be discovered what biological role, if any, these interactions play in modulating the interaction of CCR5-binding HIV strains with cell surface HS. Given the role of CCR5-binding HIV strains to HIV infection and the emerging promise of anionic polysaccharides and polysaccharide mimics towards antiviral treatment [57], a more thorough understanding of the role of cell surface GAGs on the adhesion and entry of JR-FL HIV could open new therapeutic avenues for exploration.

## Acknowledgements

This research was supported by the National Institute of General Medical Sciences (R01GM096049A to J.S.S., U01 CA207824 to R.J.W., and P41GM103390). J.S.S. discloses a significant ownership share of GenNext Technologies, Inc., a small business involved in the development of HRPF instrumentation.

